# Breeding pig transport drives the dispersal of swine influenza A virus across Europe

**DOI:** 10.64898/2026.06.01.729471

**Authors:** Marina Meester, Bram Vrancken, Fabiana Gámbaro, Simon Dellicour

## Abstract

Pigs serve as reservoirs of former human influenza A virus (IAV) H1N1 and H3N2 lineages and act as mixing vessels for diverse strains, facilitating the emergence of novel IAVs. Understanding the spread and evolution of swine IAVs (swIAVs) is therefore crucial to assess the risk of strains with zoonotic potential emerging. This study uses a phylogeographic framework to investigate the predictors of swIAV dispersal across Europe. All publicly available swIAV genomic sequences were retrieved and subsampled for the ten largest European pig-producing countries. Discrete phylogeographic reconstructions were conducted for H1, H3, N1, N2 encoding genes and all internal gene segments. Our analyses indicate that viral dispersal predominantly occurred from north-western to southern and eastern Europe, with frequent long-distance transitions between non-adjacent countries. We also extended the discrete phylogeographical analyses with generalized linear models to test the association between viral movement and potential predictors, such as live pig trade, pork trade, pig densities, farm sizes, or the geographic distance between key pig production zones. We find that breeding pig trade is the only consistently well-supported predictor of between-country transition events, whereas pork trade and geographic distance were not supported. This highlights that farms importing breeding pigs from multiple countries could act as hotspots for reassortment of diverse swIAV strains. Strengthening external biosecurity on farms with emphasis on quarantining breeding pigs, limiting long-distance transport, and implementing a One Health surveillance system for earlier detection of emerging strains, could help curb the rapid spread and evolution of swIAV in Europe.

**Author summary:** Influenza A viruses that originally came from humans are now circulating in pigs. This is a public health concern because these viruses continue to evolve in pigs and may eventually return to humans in new forms that could cause more severe disease. To better prevent this risk, we need to understand the dispersal dynamics of influenza viruses infecting pigs. For this purpose, we retrieved all publicly available European swine influenza A virus genomic sequences. Using this genomic dataset, we reconstructed how the virus has spread across Europe and investigated the external factors that may have driven the spread. Our results indicate that swine influenza A viruses frequently spread between European countries, in particular from north-western to southern and eastern Europe, and that the trade of pigs intended for breeding is strongly associated with these patterns of spread. Consequently, measures such as quarantining of breeding pigs, could help preventing swine influenza variants from spreading between and becoming established in new regions. We also find major gaps in available genomic data from several European countries. We therefore recommend stronger European surveillance programs that include monitoring influenza viruses in both animals and humans to improve early detection of new emerging viral variants.

## Introduction

Influenza A viruses (IAVs) are the most frequent cause of global pandemics in human history (1, 2). They are major respiratory pathogens in humans and many other animal species (3). IAVs transmissible between humans and other vertebrates, i.e., zoonotic IAVs, have been ranked as top priority for zoonotic disease preparedness and surveillance by experts from multiple health authorities (4, 5). This highlights the urgent need for improved understanding of the transmission and control of zoonotic IAVs. The recent epidemic of avian H5N1 IAV in over a thousand dairy cattle herds in the United States, accompanied by spillover infections to humans (6), further underscores this need. With their rapid spread, high mutation rate, and ability for interspecies transmission, IAVs represent a major One Health challenge.

Swine are considered high-risk hosts of zoonotic IAVs, because human seasonal lineages have sustained in swine populations and swine carry receptors that both human and avian influenza strains can bind to (7). This dual susceptibility allows infection with diverse IAV strains, creating optimal conditions for viral reassortment. The risk of reassortment further increases with the probability of coinfection, which in turn increases with (i) cocirculation of distinct strains in a population, and (ii) high prevalence (8) – conditions that are both present in pig populations. Namely, in one year of surveillance across fifteen European countries, at least 31 distinct swine IAV (swIAV) variants were detected, including twelve unique combinations of the surface proteins haemagglutinin (H; subtypes H1 and H3) and neuraminidase (N; subtypes N1 and N2) (9). In conventional swine farms, the high turnover of piglets and large farm sizes facilitate continuous swIAV circulation as immune pigs are replaced by susceptible ones at a rate that allows swIAVs to be maintained in the population (10). The within-farm prevalence of swIAV was estimated around 30% and the virus was found on more than half of surveilled farms in a European prevalence study (9). Together, these factors increase the likelihood that reassortants arise in pigs.

These findings substantiate the importance of understanding where swIAV variants are introduced from and what promotes the introduction of new variants into swine populations, knowledge that is crucial for designing effective swIAV control and surveillance systems. Such investigations can be carried out through phylogeographic analyses, which integrate genomic sequence data and spatial data to study the dispersal of pathogens in space and time. In particular, discrete trait phylogeographic analysis extended with a generalized linear model (GLM) can be used to evaluate the association between viral dispersal and potential predictors (11).

In this study, we aim to assess the dispersal history of swIAV across European countries, deploying Bayesian phylogeographic methods. In addition, we aim to investigate to what extent pig and pork trade, swine population demographics, and distances between countries, are associated with its dispersal dynamics. Our results indicate that the magnitude of live pig trade is the strongest predictor of swIAV dispersal between countries. By disentangling three categories of live pigs, we further distinguish which age category of pigs has the highest risk of introducing a swIAV strain into a country. Overall, the results presented in this study help identify priority-areas for the development of One Health surveillance programs targeting zoonotic IAVs.

## Results

Using sequences and associated metadata for all eight IAV gene segments retrieved from the GenBank (12) and GISAID (13) databases, we investigated the dispersal history of swIAV strains across European countries. Data were compiled separately for the H1, H3, N1, and N2 subtypes of the HA and NA gene segments, resulting in ten distinct sequence alignments. All sequences collected until August 2024 were included, and the earliest available sequence dated back to 1937. The number of sequences obtained varied between 590 (H3) and 6400 (H1) (see Table S1 for further detail on collected sequences per gene segment). Sequences were subsequently aligned, discarding sequences that were less than 90% complete, or had missing metadata such as a precise collection date or country of origin (the number of discarded sequences is detailed in Table S1). Maximum-likelihood phylogenetic trees that were first inferred to confirm a temporal evolutionary signal showed extensive intermixing of strains sampled from sixteen European countries, which warranted further phylogeographic investigations. To this end, we employed the discrete phylogeographic approach implemented in the Bayesian software package BEAST X to study the dispersal history of swIAV strains in Europe (14, 15). To overcome sampling biases across countries, we randomly downsampled the collected sequences in proportion to the estimated swIAV circulation rate in the respective countries of origin. Countries with less than ten available sequences were excluded from the analyses, resulting in a subsample from ten European countries, namely Austria, Belgium, Denmark, France, Germany, Italy, the Netherlands, Poland, the United Kingdom, and Spain, for eight out of ten gene segments. For the matrix protein (MP) gene segment, sequences were available from the Czech Republic as well, yielding eleven countries for analysis. For the H3 gene segment, only seven countries could be included, namely Belgium, Denmark, Germany, Italy, the Netherlands, Poland, the United Kingdom, and Spain. This gave a total of 132 to 919 subsampled sequences per gene segment (Table S1).

Our phylogeographic analysis revealed a notable number (>15) of well-supported transition events for every segment analysis, from Denmark to Germany, and from Germany to the Netherlands and Spain (Fig. 1) (16). For Spain, Italy, Austria and Poland, we only found strong support (BF >20) for transition events into these countries, indicating that swIAV primarily spread from other countries towards them. On the other hand, for Denmark and Germany we only found strong support for transition events from those countries and not towards. Strongly supported bidirectional spread was seen between the Netherlands and Belgium. Overall, Denmark, Germany and Spain were associated with the highest number of inferred intra-country transition events, suggesting more sustained swIAV circulation within their territory (Fig. 1).

**Figure 1.**
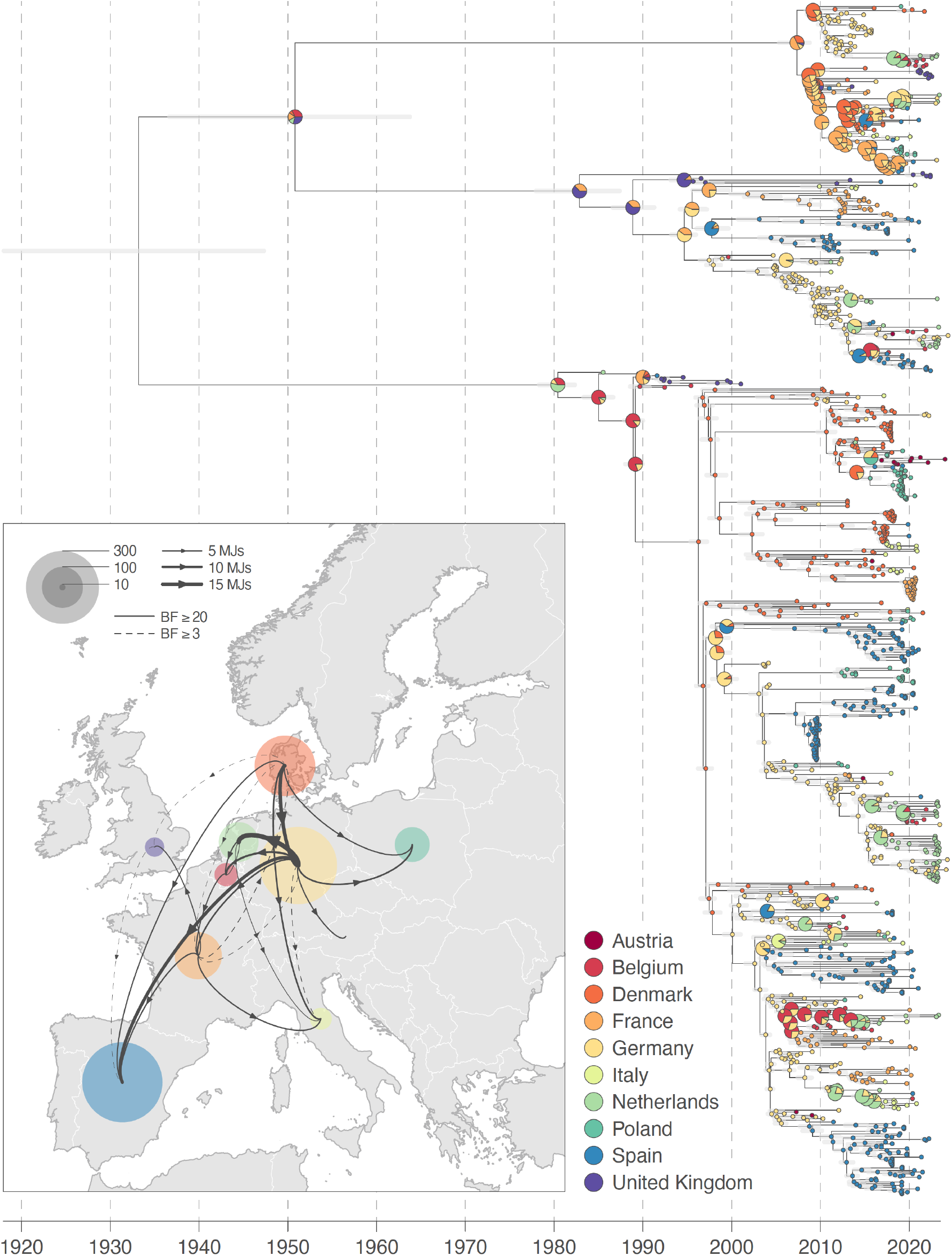
Phylogeographic reconstruction of the dispersal history of swIAV strains in Europe. Specifically, we here report the maximum clade credibility tree of the discrete phylogeographic reconstruction based on the analysis of the HA segment and conducted among ten European countries. On this tree, horizontal transparent gray line segments are associated with each internal node age estimate and reflect the 95% highest posterior density (HPD). Tip and internal nodes are colored according to the sampling and inferred location, respectively. For the internal nodes, we use a pie chart to display the posterior probabilities associated with inferred locations with a posterior probability >0.05, but only when there is not a single location inferred with a posterior probability >0.95. On the map, we then map well-supported Markov jumps (MJs) inferred by the discrete phylogeographic inference, with sampling locations displayed by dots colored by country and with a size being proportional to the number of inferred transition events within the considered country. We only report Markov jumps associated with a Bayes factor (BF) support of at least 3 and highlighted those associated with a BF support of at least 20, which can be interpreted as positive and strong supports respectively (16).

To investigate the predictors of viral transition frequencies between the included countries, we further used the GLM extension associated with the discrete phylogeographic approach (11, 14). This GLM approach returns an inclusion probability and a β coefficient per tested variable. The variables investigated were trade of live pigs, trade of offal (organs from pigs for consumption), number of pigs and farm size in the country of origin and destination, as well as the distance between the key pig production zones (centroids) of two countries.

With an inclusion probability of 0.97 to 1.00 (BF >999) for all gene segment analyses, trade of live pigs was the only consistently very strongly supported predictor of swIAV strain transition events between European countries. The corresponding β coefficients were persistently positive, indicating that higher imports of live pigs from a given country were associated with more swIAV transitions from that country (Fig. 2). We also found strong statistical support (BF >20) for the inclusion of farm size in the country of origin in five out of the ten segment-specific GLM analyses, and for inclusion of pig population size for one segment. On the other hand, distance between swine production centroids and offal trade were not statistically supported as predictors of swIAV transition frequencies between European countries in any of the analyses (Fig. 2).

**Figure 2.**
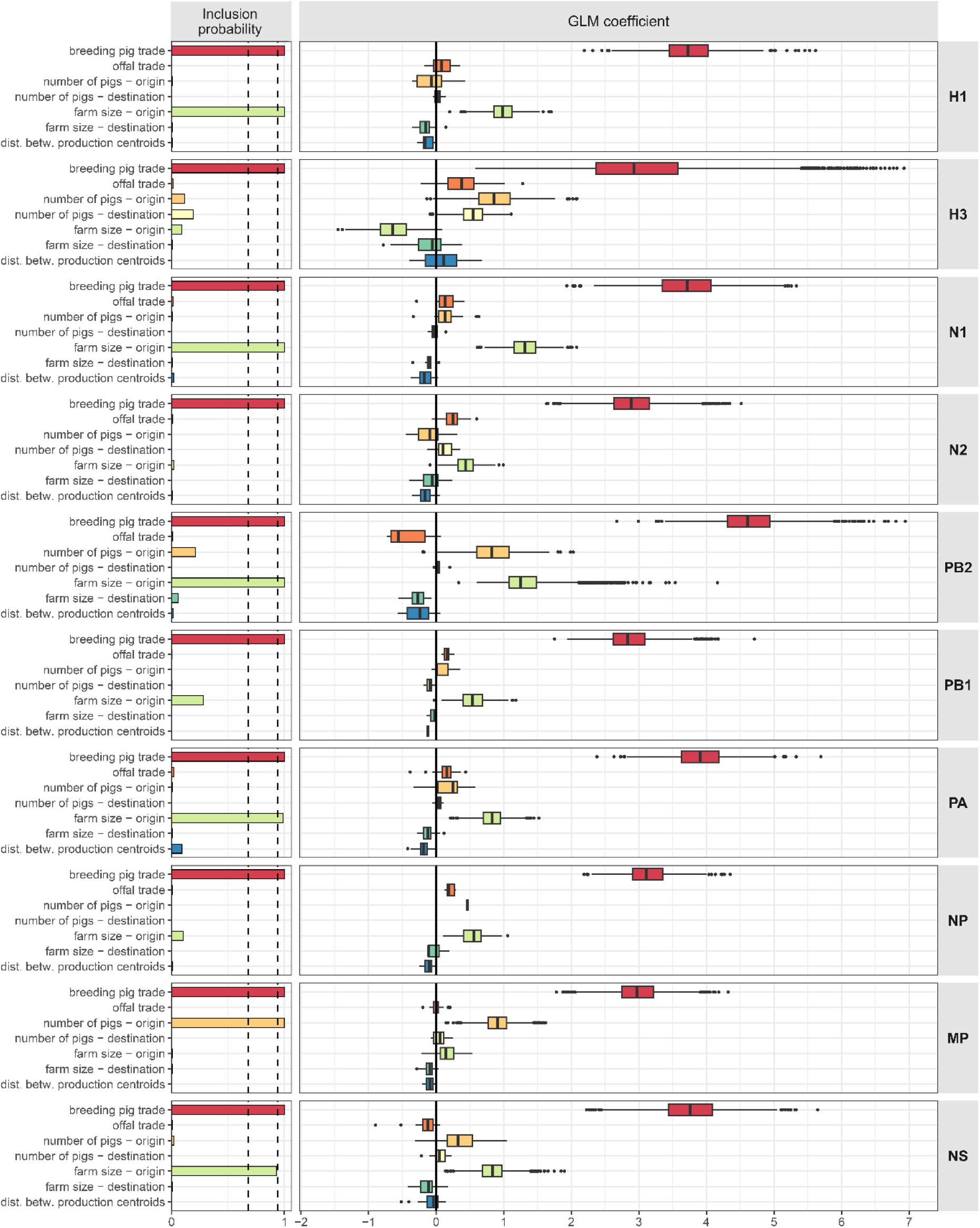
Analyses of the predictors associated with the transition frequency of swIAV among ten European countries. We here display bar plots showing the proportion of MCMC samples where the predictor was included in each segment-specific GLM analysis coupled with a discrete phylogeographic reconstruction, indicating the inclusion support of the predictor, with the two dashed lines at inclusion probabilities of 0.68 and 0.94 representing strong (Bayes factor > 20) and very strong support (Bayes factor > 150). The accompanying boxplots show the contribution of each predictor in each segment-specific GLM analysis, i.e. the GLM coefficients on a log-scale conditional on the predictor being included in the model. Dist. betw. production centroids stands for the great-circle distances between the central points of swine production in European countries.

Pigs are traded for different purposes, namely for breeding, fattening, or slaughter. International trade statistics distinguish three categories of pigs: (i) breeding (male and female pigs of different weights meant for breeding), (ii) below 50 kg (usually with the purpose to be fattened on another farm), and (iii) above 50 kg (usually meant for slaughter) (17). The traded volumes between countries were significantly correlated with each other (Spearman’s rank correlation ρ’s of 0.77, 0.80 and 0.86, p-values < 0.05). Therefore, the final GLM analysis (with results presented in Fig. 2) only includes one type of live pig trade, namely breeding pigs. However, two different approaches were conducted to distinguish which category of live pig trade best explains the swIAV dispersal history. Firstly, we included all three types of trade in the GLM analysis for four gene segments (H1, N1, N2, and PB1). In those four analyses, breeding pig trade consistently had an inclusion probability between 0.75 and 1, trade of pigs below 50 kg had an inclusion probability of 0.48 and 0.87 in two out of four analyses, but was near zero in the other two, and trade of pigs above 50 kg always had an inclusion probability near zero (Fig. S1). In addition, the GLM coefficient for breeding pigs was consistently higher than those for the other types of live pig trade (Fig. S1). Secondly, for each segment, we conducted three distinct GLM analyses, each time considering one of three pig trade variables. These analyses also revealed that breeding pig trade was consistently identified as the variable with inclusion probability near 1, and with the highest GLM coefficient, in all four segment-specific GLM analyses (H1, N1, N2, and PB1). We still found some statistical support for the inclusion of trade of pigs below and above 50 kg, but again with lower GLM coefficients than breeding pig trade. Thus, trade of breeding pigs can explain the transition events between European countries to a greater extent than trade of other pigs.

## Discussion

Zoonotic IAVs are top-priority viruses for surveillance and control plans because of their rapid evolution and wide dispersal across hosts and space (4, 5). Swine populations harbor many diverse IAV lineages, posing a risk of new variants with pandemic potential arising (9, 18). In this context, it is important to understand the factors that may favor viral spread among swine populations, as such transmission events could promote reassortment and emergence of zoonotic strains. In the present study, we investigated the direction and predictors of swIAV spread across European countries.

In 2021, more than 31 million pigs were traded between member states of the European Union (19) and we here show that this live pig trade between European countries has likely been a major driver of international swIAV spread over the last decades. SwIAV transitions were particularly driven by trade in breeding pigs and, to a lesser extent, pigs under 50 kg, but unlikely by trade in pigs above 50 kg. Studies have shown that finishing pigs (above 50 kg) are less frequently infected with swIAV than breeding and young pigs (20, 21), and are usually imported for slaughter purposes. Finishing pigs are therefore less likely than the other pig categories to come into contact with pigs in the destination country and thus to spread the virus to other farms, which confirms our finding that trade of those pigs is not relevant for spread across countries. However, our finding that trade of breeding pigs, as opposed to pigs under 50 kg, was the main driver of swIAV spread, was at first sight unexpected. Pigs under 50 kg namely have the highest prevalence of swIAV according to field studies from many countries (20) and they are traded in higher numbers than breeding pigs. Our results allow us to hypothesize that trade of breeding pigs is more related to swIAV spread due to the following reasons: commercial pig production is structured hierarchically, resembling a pyramid in which a relatively small number of breeding herds at the top supply the replacement stock to a larger number of downstream multiplier herds. Piglets from these multipliers are typically raised to intermediate weights (e.g., 25–40 kg), after which they are transported to finishing farms where they remain until slaughter weight (>100 kg) (22, 23). If a breeding herd at the top of the pyramid is infected with swIAV, and if many multiplier farmers in other countries buy its breeding pigs for restocking purposes, the virus then has multiple opportunities for establishment on destination farms. On the other hand, pigs weighing under 50 kg are typically imported as one large group from one farm to another. While this could trigger a large swIAV outbreak infecting most pigs in the destination farm, rapid depletion of susceptible pigs may ensure that the virus is unlikely to persist in the population, limiting the establishment of foreign strains in the farm. This hypothesis has been supported by a mathematical model showing that swIAV persistence after an outbreak is unlikely on farms with fewer than approximately 3,000 pigs (10). To summarize, while traded pigs under 50 kg may more frequently carry swIAV strains than breeding pigs, these strains appear less likely to establish in farms in destination countries compared with those carried by breeding pigs.

Our phylogeographic analyses reveal a swIAV dispersal pattern with notable transition events from north-western to southern and eastern Europe, with Spain and Italy having the highest number of introduction events from different countries (Fig. 1). Such a dispersal pattern is in line with the fact that Spain has increased its swine population by 30% between 2014 and 2023, while almost all other European countries either kept the same population or decreased it (24). Furthermore, the Spanish pig production sector has suffered from outbreaks of a highly pathogenic Porcine Reproductive and Respiratory Syndrome virus strain (Rosalia) since 2020, leading to an increased import of pigs (25). As for Italy, the country has a history of importing pigs because the breeding farms in the country cannot supply sufficient piglets for the demand of their traditional pork products such as hams (26). For both countries, the high demands for live pigs from abroad may have led to a high risk of importing swIAV strains. We should however interpret the dispersal patterns carefully, as discrete phylogeographic reconstructions can only infer viral transition events between sampled locations. For many EU countries, no or insufficient swIAV genomic data were available, stressing the need for increased sampling and sequencing efforts. Nevertheless, the ten countries included in the analyses correspond to the countries with largest European pig populations (24), resulting in a selection of countries that are representative for around 85% of Europe’s pig sector (24).

To mitigate the effect of sampling bias on the discrete phylogeographic analyses, we subsampled sequences based on the expected rate of a proxy for swIAV circulation rate in each considered country. This proxy was here obtained by multiplying the farm-level prevalences per country reported for 2020 (9) by the total number of pigs in each country in that year (24). For countries not included in the study of Henritzi and colleagues (9), farm-level prevalence was estimated using linear regression. Although the resulting proxy does not measure the actual number of cases, all prevalence data originated from the same study (Henritzi et al, 2020), ensuring methodological consistency. Importantly, the estimated viral circulation rates were used solely to determine the relative number of sequences retained per country. Absolute values were not critical; only the relative differences between countries were used to proportionally represent the variation in swIAV occurrence and pig population size. Given that this subsampling procedure aimed to align the resulting sampling effort with the degree of circulation, we have deliberately not incorporated the sample size as predictor in the GLM analyses, which is frequently done to assess the impact of such sampling heterogeneity when not informed based on prior pathogen occurrence knowledge (e.g. (27, 28)).

We performed an analysis for all IAV gene segments separately, because gene segments can have independent evolutionary and dispersal histories due to reassortment. By importing pigs – and thus swIAV strains – from different countries, the subtype diversity within a pig population may increase, thus increasing the risk of reassortment. Recently, methods have been published that incorporate reassortment in phylodynamic analyses [17](29). However, these methods cannot yet be integrated with the GLM analyses conducted here. Further advancements are needed to quantify the when and where of swIAV reassortment events, as well as their associated risk factors both at the country- and farm-level.

To prevent swIAV strains from crossing borders through breeding pig trade, farmers could quarantine imported pigs before bringing them into contact with the rest of the herd. The European Animal Health Law states that appropriate biosecurity measures should be taken to minimize risk of disease spread (regulation (EU) 2016/429), yet only 10 out of 24 European countries recently stated that isolation of newly introduced breeding pigs in quarantine stables is mandatory by national law (Biebaut et al, 2025). Besides, a recent survey conducted at 231 pig farms from nine European countries found that less than half of the breeding farms had a quarantine stable present (Dubbert et al, 2024). In addition, Casal and colleagues showed that although half of 172 interviewed Spanish farmers in 2007 had a separate quarantine unit available, only 20% of farmers found quarantining important (Casal et al, 2007). The limited application of quarantining may explain the high number of swIAV transition events found between countries.

Besides quarantining pigs on the importing farms, quarantining (in combination with testing) could be performed at borders. This procedure is already listed as an optional preventative measure in the Terrestrial Animal Health code by the World Organisation for Animal Health (WOAH) since 1971 (30), yet swIAV is not a WOAH-listed pathogen, making the procedure seem unnecessary in this instance. Based on our results of the spread of swIAV across Europe, we recommend the European Union to set up a pan-European surveillance system for swIAV, following One Health principles (5). This could entail increased testing of imported pigs, risk-based sampling of farms that import pigs from various other countries, or farms that house multiple animal species, and sampling of professionals with flu symptoms shortly after contact with pigs. Environmental air samples in areas where pig farms are located in residential areas may be considered as well. Finally, scaling down live pig trade across Europe will not only reduce the long-distance spread of swIAV strains on the continent, but also improve animal welfare of pigs and reduce the environmental burden of pig production. Therefore, this is a crucial measure to undertake.

## Methods

### Sequence data

#### Selection and processing of RNA sequences

SwIAV genomic sequences were retrieved from the GenBank (12) and GISAID (13) public databases. All uploaded sequences from the eight IAV gene segments uploaded until August 1, 2024, with swine as host species and Europe as geographic region were downloaded along with associated metadata (i.e. the country of origin and collection date). Duplicate sequences from the two databases were filtered out using the identity of ‘GenBank_Title’ in NCBI and ‘Isolate_Name’ in GISAID metadata annotations.

Sequences were aligned per gene segment, using the FFT-NS-2 algorithm implemented in the program MAFFT v7.520 (31). The resulting alignments were then converted to tibbles using the function “as_tibble.bioseq_dna” of the “bioseq” R package (v4.3.3) in RStudio (v2023.12.1.402) (32) to remove nucleotide positions with over 60% gaps (33). Subsequently, sequences with more than 10% of gaps were removed from the dataset. In addition, sequences for which only the year of collection was reported were excluded using the “tidyverse” R package (34), except for those older than 2004 due to scarcity of old sequences. Recombination within gene segments was assessed using the pairwise homoplasy index (PHI) as implemented in the program SplitsTree (35).

#### Phylogenetic trees and temporal signal

Maximum-likelihood (ML) phylogenetic trees were constructed with RaxML-NG with an HKY+Γ substitution model for all gene segments, and branches were colored by country in IcyTree to identify whether monophyletic clades are shared by more than one country (36). Temporal signal was assessed by root-to-tip regressions on the ML trees using the program TempEst (37). Outliers were removed based on residuals higher than 0.03 until a coefficient of determination (R^2^) of at least 0.6 was found. The number of outliers per gene segment can be found in Table S1. The HA gene was subtyped using BV-BRC’s subspecies classification tool (38).

### Subsampling procedure

Spatial and temporal sampling heterogeneity are known to potentially bias discrete phylogeographic reconstructions (14, 39, 40), with a tendency to overestimate transitions of viral strains from the most sampled locations. To circumvent the impact of sampling bias, one approach consists in downsampling the datasets relative to the circulation rate of the pathogen approximated in each sampled location. For swIAV, Henritzi and colleagues (2020) determined the farm-level prevalence in European countries based on one year of surveillance (9). To approximate the relative circulation rate of swIAV in pigs in each considered country, we assumed that this is relative to the farm-level prevalence multiplied by the number of pigs present for each country. To derive a proxy of the farm-level prevalence for countries for which prevalence was not estimated before, we fitted a multivariable linear model with farm-level prevalence as the dependent variable, and tested the number of pigs (24), number of pig farms (41), farm size, gross domestic product (42), and country size in km^2^ (43) as independent variables. The best performing linear model, using Akaike’s Information Criterion, was:

Farm-level prevalence = β_0_ + β_1_·farm size + β_2_·number of farms + β_3_·(farm size · number of farms) + ε, with an adjusted coefficient of determination (R^2^) of 0.53 and an associated p-value < 0.05. Using this linear model, the farm-level prevalence was predicted for the remaining countries. Finally, the prevalence was multiplied by the number of pigs present per country in one year to obtain a proxy for the swIAV circulation rate in each country.

The downsampling of sequences was thus guided by the swIAV circulation rate proxy and aimed at achieving a number of sequences proportional to the actual level of viral circulation in each country. Downsampling was performed randomly within each country and separately for each gene segment. When the number of available sequences in a country was lower than the target based on the corresponding proxy value, fewer sequences were included from other countries to maintain a proportional representation. Countries with fewer than ten available sequences were excluded from the analysis. If, after downsampling, a country had fewer than ten sequences remaining, it was upsampled (e.g., by resampling without replacement) to ensure a minimum of ten sequences per country.

### Discrete phylogeographic analyses

We conducted the phylogeographic analyses using the discrete diffusion model (14) implemented in the software package BEAST X v10.5.0 (44) coupled with the BEAGLE v4.0.0 library for improved computational efficiency (45). In those analyses, we modeled the asymmetric transitions between countries using a continuous-time Markov chain (CTMC) framework, as described by (14). In this approach, movement among *K* discrete locations – i.e. European countries in the present study – are captured by a *K*×*K* infinitesimal rate matrix *Λ*. Each element *Λ*_*ij*_ represents the instantaneous and relative transition rate from location *i* to *j*, for which we also obtain a Bayes factor (BF) support. To study the relative contribution of potential predictors to swIAV dispersal among countries, we then investigated predictors for dispersal using the GLM extension (11) of the discrete phylogeographic model; again conducting a distinct analysis per segment. With these GLM analyses, we obtained, for each predictor, a regression coefficient as well as an inclusion frequency and associated BF support (11). We assume a BF higher than 3 indicates positive support for a predictor’s contribution, higher than 20 strong support and higher than 150 indicates very strong support, as proposed by Kass and Raftery (16).

#### Predictors of transition frequencies between countries

We assessed four categories of predictors of transition frequencies between countries: live pig trade, pork trade, pig population, and distance-based predictors. For live pig trade, public database BACI (17) provided yearly data on import and export of pigs with a weight below 50 kg, above 50 kg, and breeding pigs, in units of total weight traded (kg). With regard to pork products, data are available for a wide range of cuts and preservation methods. IAVs generally do not infect muscle tissue yet primarily infect respiratory tissues (46). Therefore, we selected the extent of offal (organs) trade between countries as one predictor. In addition, meat (muscle tissue) may be contaminated due to respiratory secretions of infectious pigs at slaughterhouses (47), so trade of the two largest pork cuts, fresh half carcasses and hams, were selected as well. With regard to the pig population, the number of pigs per year, number of farms, average number of pigs per farm, and number of pigs per km^2^ were considered and collected from Eurostat (24, 41). Distance-related parameters were defined as follows: having shared borders; great-circle distance between the country’s centroids, and great-circle distance between the centroids of swine production in each country. A centroid of swine production was defined as the average geographic location of pigs per country, weighted by the number of pigs per 10 km^2^ grid, obtained from the GLW4 database (48, 49). Metadata were reported for different time periods across parameters. For example, the number of farms per country was available for 2010, 2013, 2016, and 2020, whereas annual data on pig populations were available from 2007 onward, and trade data from 1995 onward. To ensure comparability across parameters, we calculated average values for the period 2010 – 2020 for use in the main analyses. As a sensitivity analysis, we repeated all analyses using only 2020 values for each predictor; this yielded consistent results in terms of inclusion probabilities and the direction of GLM coefficients (results not shown).

Correlation between predictors was assessed in R prior to the GLM analyses. For dissimilarity matrices (i.e., trade of pigs and pork between countries), Spearman’s rank correlation coefficients (*ρ*) were calculated using the mantel function of the “vegan” R package V2.7-2 (50). For pig population data (i.e., average farm size and pig density) they were calculated using the “cor.test” (“method = spearman”). In case predictors were significantly correlated (p-value < 0.05) only the potential predictor that made most biological sense was included in downstream analyses, resulting in the selection of breeding pig trade, offal trade, as offal trade was correlated with trade of the other pork cuts, number of pigs present, average farm size, and distance between swine production centroids In the main analyses. Because we were interested in which category of live pig trade was most important for swIAV dispersal, but the trades of the three categories of live pigs were significantly correlated, we performed additional analyses for four gene segments, H1, N1, N2 and PB1. In the first approach, we included all three categories of live pig trade in one analysis. In the second approach, each category of live pig trade was included in a separate analysis, keeping the other predictors constant, leading to a total of twenty additional analyses. For the pig population parameters, we specified an origin and destination predictor value for each pairwise transition rate, giving a total of seven predictors. We added pseudocounts for predictors that had zero entries and subsequently performed log-transformation and standardization on the predictor data prior to the GLM analyses.

#### Model assumptions

In those phylogeographic analyses, swIAV evolution was modeled using an HKY substitution model with Γ-distributed rate variation across four categories, allowing for empirical estimation of nucleotide frequencies (51). For the tree prior, we used the nonparametric skygrid coalescent model (52) and employed an uncorrelated relaxed clock model with an underlying lognormal distribution, as the 95% highest posterior density interval (HPDI) of the coefficient of variation in preliminary analyses did not include 0. For sequences with incomplete sampling dates, tip-date sampling from a uniform prior that spans the entire month or year was performed (53). MCMC analyses were run with 200 to 800 million iterations to achieve adequate statistical mixing and effective sample sizes (ESS) > 200 on all relevant parameters, as assessed with the program Tracer v1.7.2 (54). Maximum clade credibility (MCC) trees were retrieved and annotated after discarding a burn-in of 10% (or more if needed) using the program TreeAnnotator (55). The visualisation of the discrete phylogeographic reconstruction was generated with a custom R script.

## Data availability statement

R scripts as well as input and output files related to the different phylogeographic analyses conducted in this study are available at https://github.com/MarinaMeester/stop_siv/

## Acknowledgments

The authors acknowledge support from the COST Action ESFLU, CA21132, which provided a travel grant enabling the first author to visit the Spatial Epidemiology lab of the ULB, Belgium.

